# Cardiac-respiratory coordination during memory encoding predicts performance

**DOI:** 10.64898/2026.01.14.699577

**Authors:** Xindan Zhang, Timo Kvamme, Yoko Nagai, Juha Silvanto

## Abstract

Cardiorespiratory coupling (i.e., the phase synchronization between cardiac and respiratory rhythms) varies with physiological state, but whether coupling strength functionally predicts cognitive performance remains unknown. Here we demonstrate that cardiorespiratory coupling predicts trial-level accuracy during the initial formation of mental representations but not during their subsequent retention, revealing temporal specificity in the relationship between bodily rhythms and cognition. We experimentally manipulated coupling via paced breathing (slow: 6 bpm vs. fast: 20 bpm) while participants performed mental imagery and working memory tasks with continuous physiological monitoring. Slow breathing enhanced parasympathetic activity (respiratory sinus arrhythmia), which strengthened baseline cardiorespiratory coupling. This coordinated autonomic state persisted into subsequent cognitive trials, predicting trial-by-trial accuracy specifically during the initial encoding phase when internal representations were actively constructed, but not during subsequent maintenance when representations were sustained. This temporal dissociation indicates that cardiorespiratory coordination selectively predicts performance when the brain constructs internal representations under elevated processing demands. Experimental manipulation of breathing improved behavioral accuracy, validating coupling’s functional relevance. These findings establish cardiorespiratory coupling as a physiological state marker that reliably predicts encoding success, providing a measurable target for interventions targeting memory and mental representation.

## 1. Introduction

The brain operates in close conjunction with the autonomic nervous system. Ascending signals from the heart, lungs, and viscera continuously shape neural processing, with cardiac and respiratory rhythms modulating cortical excitability, attention, and memory (Park et al., 2014; Zelano et al., 2016; Garfinkel et al., 2014). Stimuli presented at specific phases of the cardiac cycle show altered cortical processing, respiratory rhythms entrain hippocampal oscillations during memory encoding, and disrupted interoceptive signaling characterizes anxiety, trauma, and neurodevelopmental conditions (Critchley & Harrison, 2013). Understanding how bodily rhythms scaffold cognition is thus fundamental to both basic neuroscience and clinical intervention.

Prior research has examined cardiac and respiratory effects on cognition separately, demonstrating that each rhythm independently modulates neural activity and behavior. Cardiac signals influence perceptual thresholds and metacognitive judgments through phasic baroreceptor input (Park et al., 2014), while respiratory oscillations entrain limbic structures, with memory performance varying across breathing phases (Zelano et al., 2016; Herrero et al., 2018). However, the heart and lungs do not function in isolation; their coordination is mediated by vagal pathways that synchronize cardiovascular and respiratory oscillators in the brainstem (Schäfer et al., 1998). During slow breathing, strong parasympathetic activation amplifies this inherent synchronization: the vagus nerve modulates heart rate tightly in phase with breathing, creating cardiorespiratory coupling that varies systematically with physiological state—strengthening during meditation and weakening under stress (Cysarz & Büssing, 2005; Zhang et al., 2010). This coupling may stabilize cerebral blood flow and optimize metabolic support when neural computations impose elevated demands (Willie et al., 2014). Yet research has treated coupling primarily as a state marker, comparing average levels between conditions. Whether coupling strength actively predicts cognitive performance at trial-level within individuals remains unknown.

Critically, if cardiorespiratory coordination influences cognition, it may do so with temporal specificity across different phases of mental processing. Cognitive operations such as working memory and mental imagery unfold in distinct phases with markedly different computational demands. Constructing internal representations requires recruiting and synchronizing distributed neural populations under elevated metabolic load, demanding stable oxygen delivery and cerebral perfusion. In contrast, maintaining already-established representations imposes minimal ongoing metabolic demands. If synchronized bodily rhythms provide physiological scaffolding for cognition, this benefit should be maximal precisely when demands peak—during active mental construction—rather than during passive retention of stabilized representations. Testing this temporal specificity is essential to understanding whether coupling functions as a domain-general state marker or as dynamic physiological support that assists cognition specifically when computational and metabolic demands are highest.

We experimentally manipulated cardiorespiratory coupling via paced breathing (slow: 6 breaths/min vs. fast: 20 breaths/min) while participants performed mental imagery and working memory tasks with continuous physiological monitoring. Slow-paced breathing robustly enhances parasympathetic tone and strengthens cardiorespiratory coupling (Laborde et al., 2022; Ren & Zhang, 2019), providing a controlled experimental tool for testing whether coupling functionally supports cognitive performance or merely reflects physiological state.

Both tasks required maintaining internal representations of identical geometric shapes but differed in whether representations were generated from verbal cues (imagery) or visually presented (working memory), allowing us to test whether coupling effects generalize across encoding modalities. We measured respiratory sinus arrhythmia (RSA), heart rate variability (RMSSD), and cardiorespiratory coupling (instantaneous phase coherence) continuously from baseline through task performance, analyzing coupling strength separately during encoding and maintenance phases. We hypothesized that coupling would predict trial-level accuracy specifically during encoding—when representations are actively constructed under high metabolic demand—not during passive maintenance, establishing it as an active physiological mechanism supporting dynamic mental operations.

## 2. Methods

### 2.1. Participants

Thirty-two participants (18 females, 14 males; mean age = 23.6 years, SD = 2.4) with normal or corrected-to-normal vision were recruited from the University of Macau. All participants provided written informed consent and received compensation upon completion. Sample size (N = 32) was based on similar within-subjects breathing manipulation studies (Luo et al., 2025; Ren & Zhang, 2019). With 48 trials per task (24 per breathing condition), the repeated-measures design provided adequate statistical power (> .80) to detect medium within-subjects effects (d ≥ 0.50) for the primary breathing comparisons. For the primary trial-level analysis testing whether coupling predicts accuracy, statistical power derives from the number of observations (2,686 trials) rather than participants, providing >95% power to detect the observed effect (OR = 1.85) and >80% power for effects as small as OR ≥ 1.30. Post-hoc sensitivity analysis confirmed adequate power for all reported effects. The study was approved by the University of Macau Research Ethics Committee. All procedures were conducted in accordance with the Declaration of Helsinki.

### 2.2. Experiment design

Participants completed two sessions separated by three days in a 2 (breathing: slow vs. fast) × 2 (task: mental imagery vs. working memory) within-subjects design (Fig. 1). Each session tested one task under both breathing conditions. Within each session, participants completed: (1) 5 minutes of paced breathing (6 bpm slow or 20 bpm fast), (2) 24 task trials, (3) 30-second rest, (4) 5 minutes of the alternative breathing rate, (5) 24 trials of the same task. Task order across sessions and breathing order within sessions were fully counterbalanced across participants.

**Figure 1.**
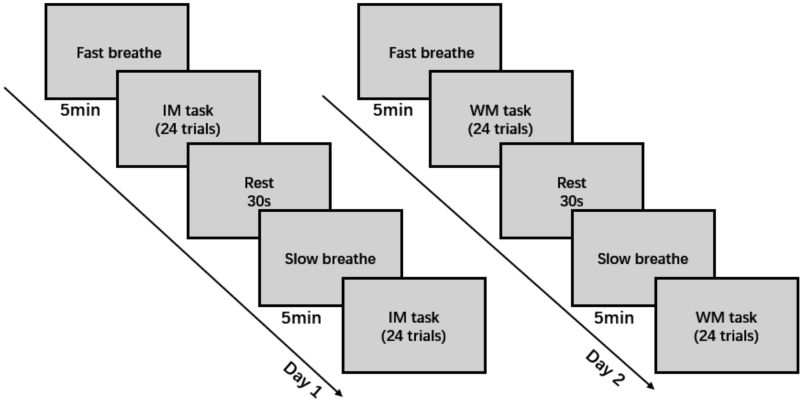
Experimental timeline. Participants completed two sessions separated by three days. Each session tested one cognitive task (mental imagery or working memory) under both breathing conditions (slow and fast). Task order across sessions and breathing order within sessions were counterbalanced across participants.

### 2.3. Imagery and working memory tasks

Tasks were adapted from Jacobs (2018) with modifications (Fig 2). In the Mental Imagery (IM) task, each trial presented: a fixation cross (1000 ms), a shape name cue—diamond, triangle, or parallelogram (1000 ms), four placeholders demarcating the imagery area (3000 ms; extended from 1500 ms based on pilot testing), and a maintenance delay (4000 ms). During the placeholder period, participants generated a vivid mental image of the cued shape; during the delay, they maintained it. A probe dot then appeared at a fixed distance from the shape’s boundaries, and participants judged whether it fell inside or outside their imagined shape. After responding, participants rated imagery vividness and judgment confidence (1-4 scales). The Working Memory (WM) task was identical except the shape was presented visually (1500 ms) followed by a noise mask (200 ms) instead of the placeholder display.

**Figure 2.**
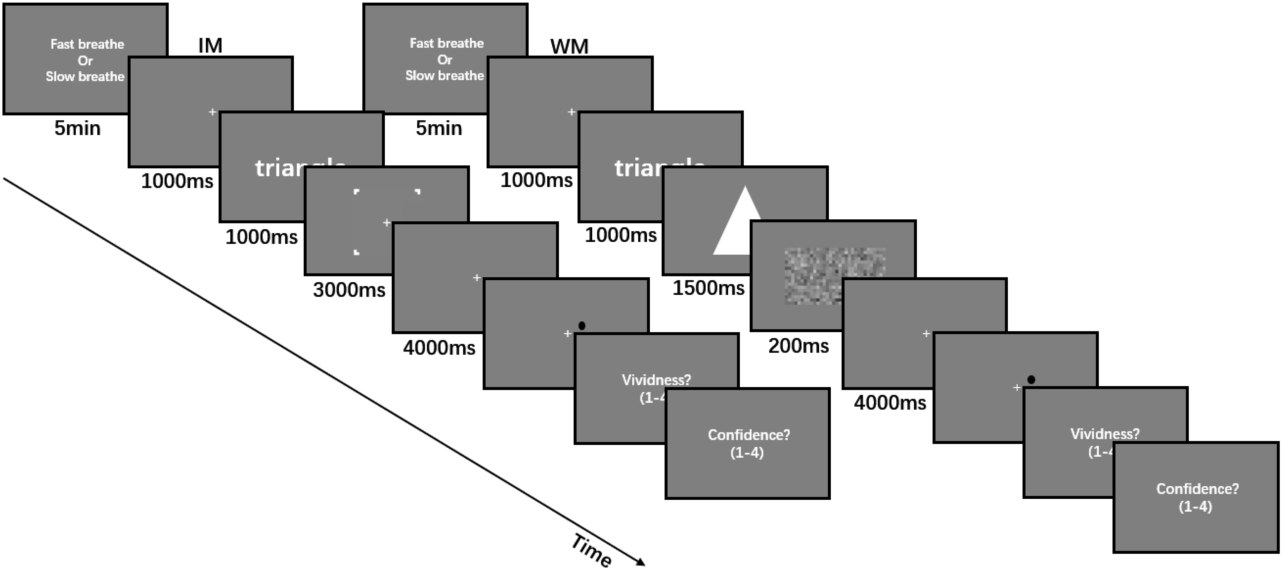
Experimental paradigms. (Left) Mental Imagery (IM) task trial structure: fixation (1000 ms), shape name cue (1000 ms), placeholder display (3000 ms) during which participants generated a vivid mental image of the cued shape, delay period (4000 ms) for image maintenance, probe dot judgment (inside/outside imagined shape boundaries), and vividness/confidence ratings (1-4 scales). (Right) Working Memory (WM) task: identical to IM except the shape was presented visually (1500 ms) followed by a noise mask (200 ms) instead of the placeholder display.

### 2.4. Electrocardiogram and Respiratory data acquisition

Electrocardiogram (ECG) and respiratory (RESP) signals were acquired at a sampling rate of 2000 Hz using a BioPac MP160 data acquisition system with AcqKnowledge software. Baseline wander was removed from the raw ECG using a zero-phase 2nd-order Butterworth high-pass filter with a 0.5 Hz cutoff. QRS complexes were enhanced using a discrete wavelet transform (Symlet-4, six decomposition levels) with selective reconstruction and weighted combination of the detail coefficients from levels 4, 5, and 6 (heuristic weights: 0.3, 1, and 1.5). R-peaks were identified via adaptive peak detection, with the minimum peak distance set to 0.35 times the sampling frequency and the minimum peak prominence set to 30% of the amplitude range between the 97th and 80th percentiles of the enhanced signal. Respiratory signals were linearly detrended and standardized (zero mean and unit variance) prior to downstream analyses.

### 2.5. Data analysis

All statistical analyses were conducted in R (version 4.4.1). We used the lme4 package (version 1.1-35.5) to fit mixed-effects models, and Satterthwaite method for degrees of freedom and p-values calculation. The significance level for all statistical tests was set at α = .05. For continuous physiological measures (e.g., RSA, RMSSD, and instantaneous phase coherence (IPC)), we employed linear mixed models estimated via restricted maximum likelihood (REML). For models where residual diagnostics indicated deviations from normality, we employed nonparametric bootstrapping (1,000 resamples) to obtain percentile confidence intervals for fixed effects, using the boot package (version 1.3.30) and robust linear mixed models via the robustlmm package (version 3.3.3), which uses an adaptive weighting scheme to down weight influential observations. For the binary behavioural measure (trial-level accuracy, correct = 1, error = 0), we used generalized linear mixed models with a binomial distribution and a logit link function. Model parameters were approximated using adaptive Gauss–Hermite quadrature, and the bobyqa optimizer was applied to ensure convergence. Both types of models included a random intercept for participants (1 | participant) to account for between-subject variability.

### 2.5.1. Trial Phase Segmentation

Cardiorespiratory coupling estimation via instantaneous phase coherence requires sufficient duration to capture respiratory oscillations, particularly during slow breathing (6 bpm = 10 seconds per cycle). Analysis windows shorter than ∼4 seconds yield unstable phase estimates. We therefore segmented each trial into two 4.5-second windows.

Mental imagery trials (∼9 seconds) were longer than working memory trials (∼7.7 seconds) because mental image construction by nature requires approximately 1-1.5 seconds longer than WM encoding. Maintenance periods were matched. For imagery, windows were non-overlapping: encoding period (0-4500 ms) captured cue presentation and active construction of the mental image; maintenance period (4500-9000 ms) captured sustained maintenance of the mental image. For working memory, the shorter encoding period necessitated ∼1.3-second overlap (encoding: 0-4500 ms; maintenance: 3200-7700 ms). Identical 4.5-second durations across tasks ensured coupling differences reflected task effects rather than window-length artifacts. The encoding window captured initial stimulus processing and consolidation into a stable representation, while the maintenance window reflected sustained retention of already-stabilized representations.

#### 2.5.2. Heart Rate Variability and Respiratory Sinus Arrhythmia Quantification

Heart rate variability was assessed using the root mean square of successive differences between normal R-R intervals (RMSSD), a time-domain index of parasympathetically mediated cardiac variability. Respiratory Sinus Arrhythmia (RSA) was quantified as the log-transformed spectral power (Welch’s method, 16 s windows, 50% overlap) of the IBI time series within the respective fixed frequency band, which is 0.07-0.13 Hz for slow breathe condition and 0.25-0.40 Hz for fast breathe condition.

#### 2.5.3. Cardiorespiratory Coupling Quantification

Cardiorespiratory coupling was quantified using instantaneous phase coherence (IPC) via the phase locking value (PLV). Instantaneous phases of heat rate and respiratory signals were extracted using the Hilbert transform. PLV was calculated as: PLV = |mean(exp(i × (φ_resp – φ_hr)))|, where φ represents the instantaneous phase. PLV ranges from 0 (no synchronization) to 1 (perfect synchronization).

Baseline cardiorespiratory coupling was quantified using the final 60 seconds of the 5-minute paced breathing period. This window was selected for two reasons. First, coupling strength requires time to stabilize following a change in breathing rate; by the final minute, participants have completed four minutes of entrainment, representing steady-state coordination rather than transient adjustment. Second, temporal proximity to the subsequent task period makes the final minute most predictive of the physiological state participants will maintain during cognitive performance.

## 3. Results

### 3.1. Cardiorespiratory Coupling Predicts Trial-Level Accuracy During encoding Phase

Cardiorespiratory coupling during the encoding phase significantly predicted trial-level accuracy across both mental imagery and working memory tasks (b = 0.615, SE = 0.215, z = 2.865, p = .004, OR = 1.85, R²m = 0.010, R²c = 0.035; Fig. 3A). In contrast, coupling during the subsequent maintenance phase showed no relationship with performance (b = 0.160, z = 0.731, p = .465, R²m = 0.005, R²c = 0.029; Fig. 3B). To formally test whether coupling’s effect on accuracy differed between encoding and maintenance phases, we fit a generalized linear mixed model including coupling, phase (encoding vs maintenance), and their interaction, with random intercepts for participants. The phase × coupling interaction was significant (b = -1.225 SE = 0.625, z = -1.961, p = .0498), confirming that coupling strength predicted accuracy during encoding but not maintenance. This temporal dissociation indicates that cardiorespiratory coordination supports cognition specifically during active mental operations, not during stable retention of established representations.

**Figure 3.**
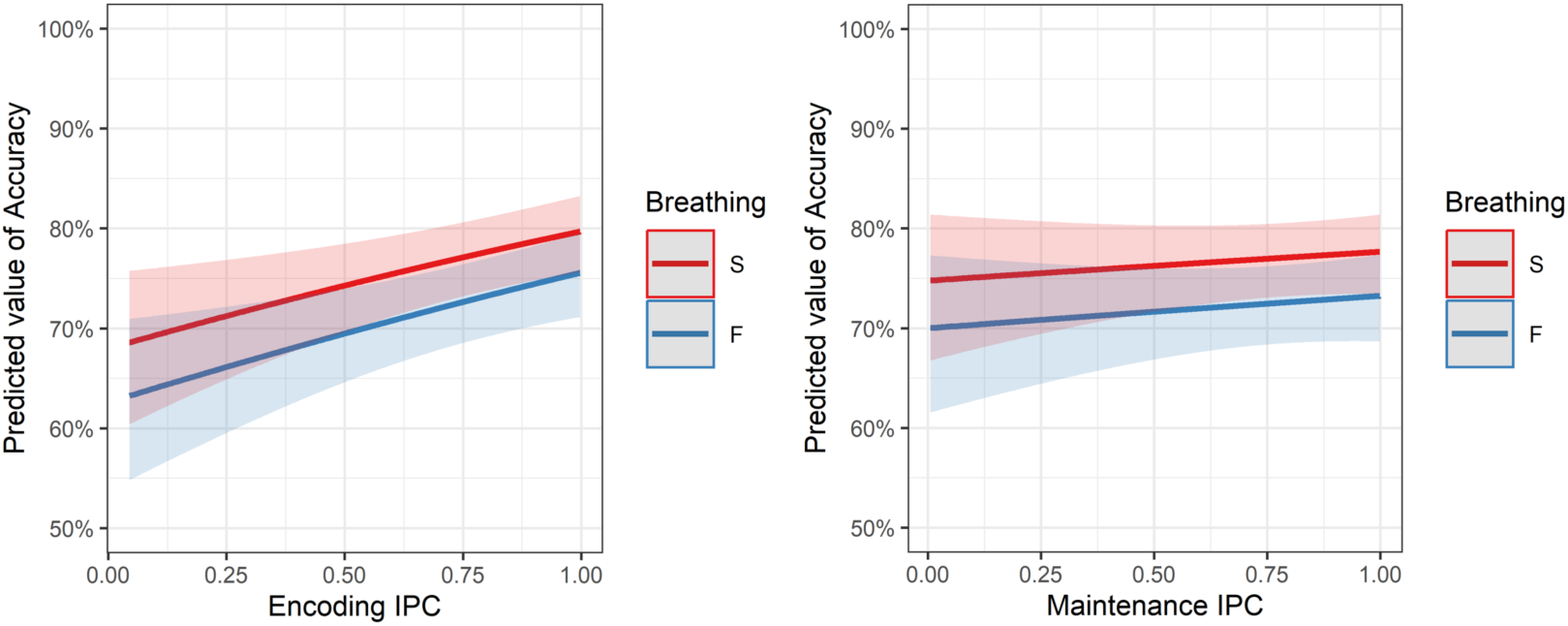
Cardiorespiratory coupling predicts accuracy during encoding but not maintenance. (A) Coupling strength during the encoding phase positively predicts trial-level accuracy (b = 0.615, p = .004), with slow breathing producing stronger coupling than fast breathing. (B) Coupling during the maintenance phase shows no relationship with accuracy (b = 0.160, p = .465). Red: slow breathing (6 bpm); Blue: fast breathing (20 bpm). Lines show predicted accuracy from logistic regression with 95% confidence intervals. IPC = instantaneous phase coherence.

This relationship remained significant when controlling for baseline parasympathetic activity (as measured during the breathing manipulation): encoding-phase coupling predicted trial-level accuracy even when including baseline RSA and RMSSD as covariates (coupling: b = 0.616, SE = 0.215, z = 2.872, p = .004; RSA: b = -0.019, SE = 0.060, z = -0.311, p = .755; RMSSD: b = 0.004, SE = 0.004, z = 0.877, p = .381). This indicates that cardiorespiratory synchronization during encoding contributes to performance independently of the parasympathetic state established during paced breathing Coupling strength did not significantly predict subjective vividness ratings during encoding (b = 0.050, SE = 0.046, t (2654) = 1.090, p = .276, 95% CI [-0.040, 0.141], d = 0.10) or maintenance (b = 0.009, SE = 0.046, t (2654) = 0.185, p = .853, 95% CI [-0.082, 0.099], d = 0.02) phases. Confidence ratings showed a marginal trend during encoding (b = 0.112, SE = 0.063, t (2660) = 1.785, p = .074, 95% CI [-0.011, 0.236], d = 0.17) but no relationship during maintenance (b = 0.027, SE = 0.063, t (2659) = 0.423, p = .673, 95% CI [-0.097, 0.151], d = 0.04). These results reveal a dissociation between objective performance and metacognitive awareness, with coupling influencing computational precision but not subjective monitoring of representation quality.

### 3.2. Experimental Manipulation of Coupling via Paced Breathing

To test whether the coupling-accuracy relationship holds under experimental manipulation, we used paced breathing to modulate cardiorespiratory coordination. Slow breathing (6 bpm) robustly enhanced parasympathetic activity compared to fast breathing (20 bpm), indexed by respiratory sinus arrhythmia (RSA: b = 1.828, SE = 0.199, t(89) = 9.205, p < .001, d = 2.34, R²m = 0.445, R²c = 0.706) and heart rate variability (RMSSD: b = 0.395, SE = 0.072, t(88) = 5.476, p < .001, d = 1.39, R²m = 0.203, R²c = 0.680; see Fig. 4).

**Figure 4.**
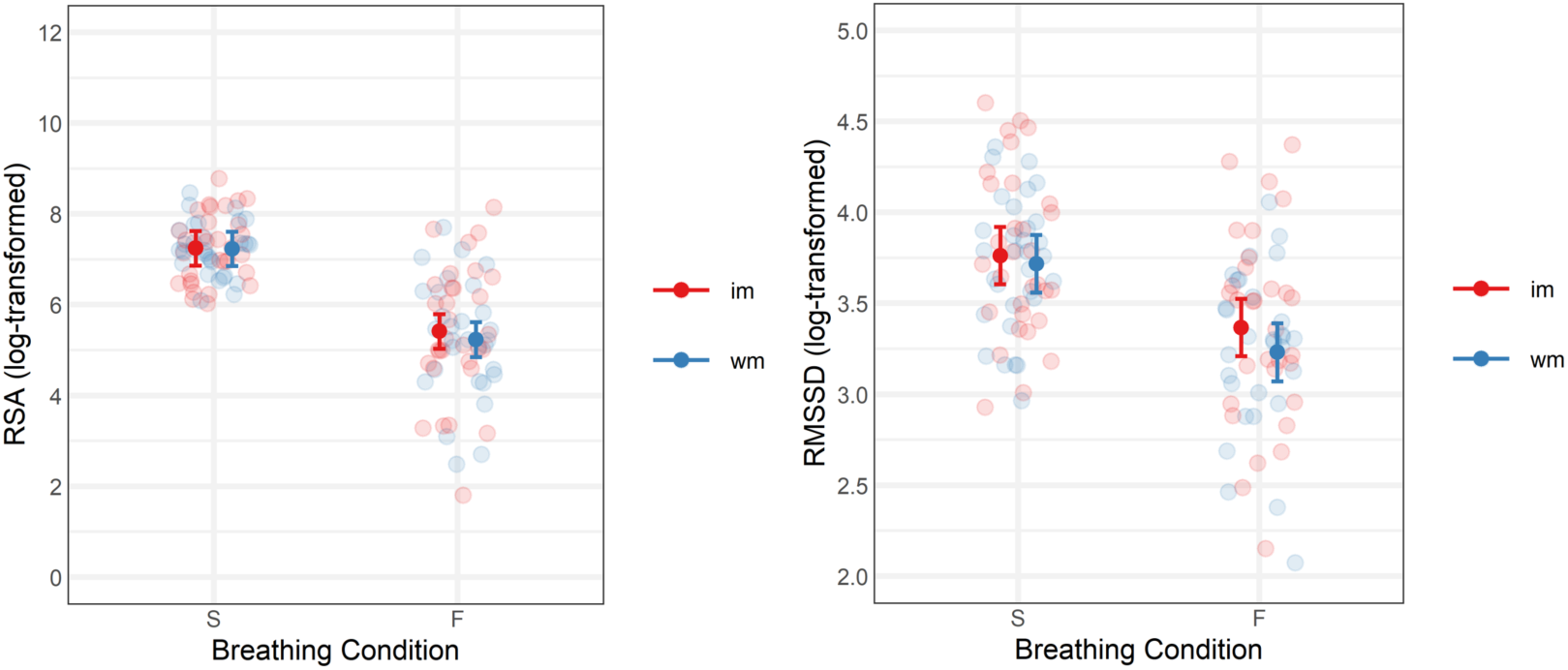
Effects of paced breathing on RSA and heart rate variability. A) Respiratory Sinus Arrhythmia (RSA) and B) Root Mean Square of Successive Differences (RMSSD), indices of parasympathetic activity, were significantly higher during the slow-paced breathing condition compared to the fast-paced breathing condition. S=Slow breathing; F= Fast breathing; im=Imagery task; wm = working memory task.

This parasympathetic enhancement facilitated cardiorespiratory coordination: RSA during the breathing period significantly predicted baseline coupling strength (b = 0.062, SE = 0.014, t(80) = 4.587, p < .001, d = 0.42, R²m = 0.652, R²c = 0.673; Fig. 5), and consequently, baseline coupling was substantially higher during slow breathing (b = 0.328, SE = 0.039, t(89) = 8.356, p < .001, d = 2.12, R²m = 0.588, R²c = 0.646).

**Figure 5.**
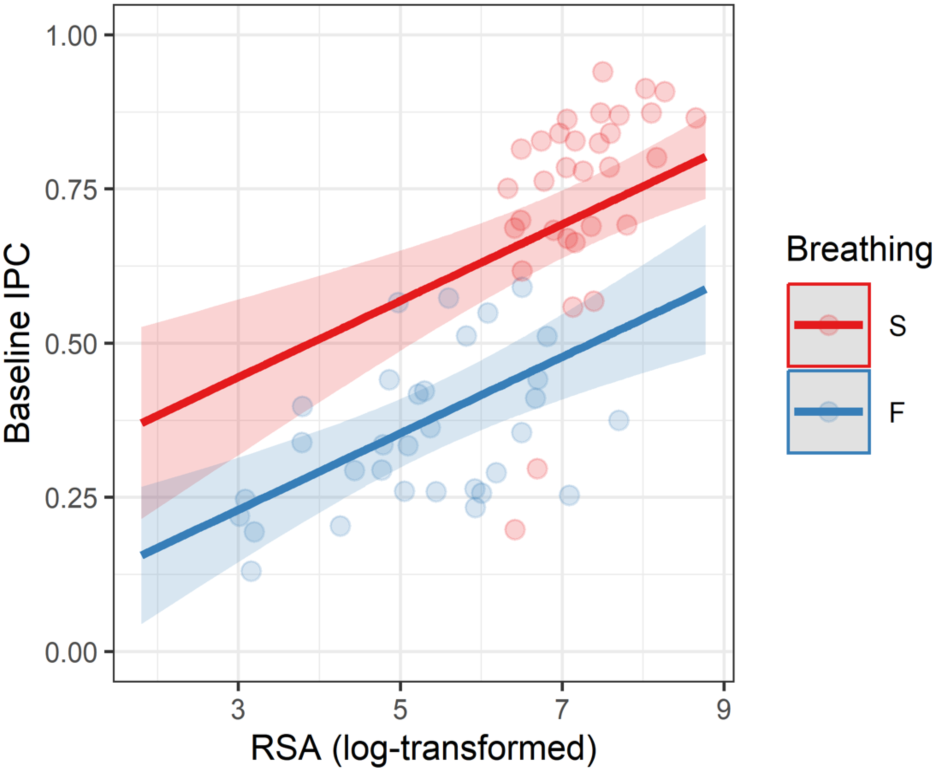
– Respiratory sinus arrhythmia predicts baseline cardiorespiratory coupling. RSA measured during the 5-minute paced breathing period positively predicted baseline coupling strength measured during the final 60 seconds of paced breathing (b = 0.062, p < .001, d = 0.42). This baseline coupling represents the coordinated autonomic state established before cognitive task performance. Points represent individual participants; red = slow breathing (6 bpm), blue = fast breathing (20 bpm). Lines show linear regression fits with 95% confidence intervals. IPC = instantaneous phase coherence.

This enhanced coupling state persisted into subsequent task performance. Stronger baseline coupling predicted stronger encoding-phase coupling (b = 0.067, SE = 0.029, t(2399) = 2.327, p = .020, d = 0.33, R²m = 0.007, R²c = 0.106; Fig. 6A), demonstrating that the coordinated autonomic state established during paced breathing carried forward into cognitive processing.

**Figure 6.**
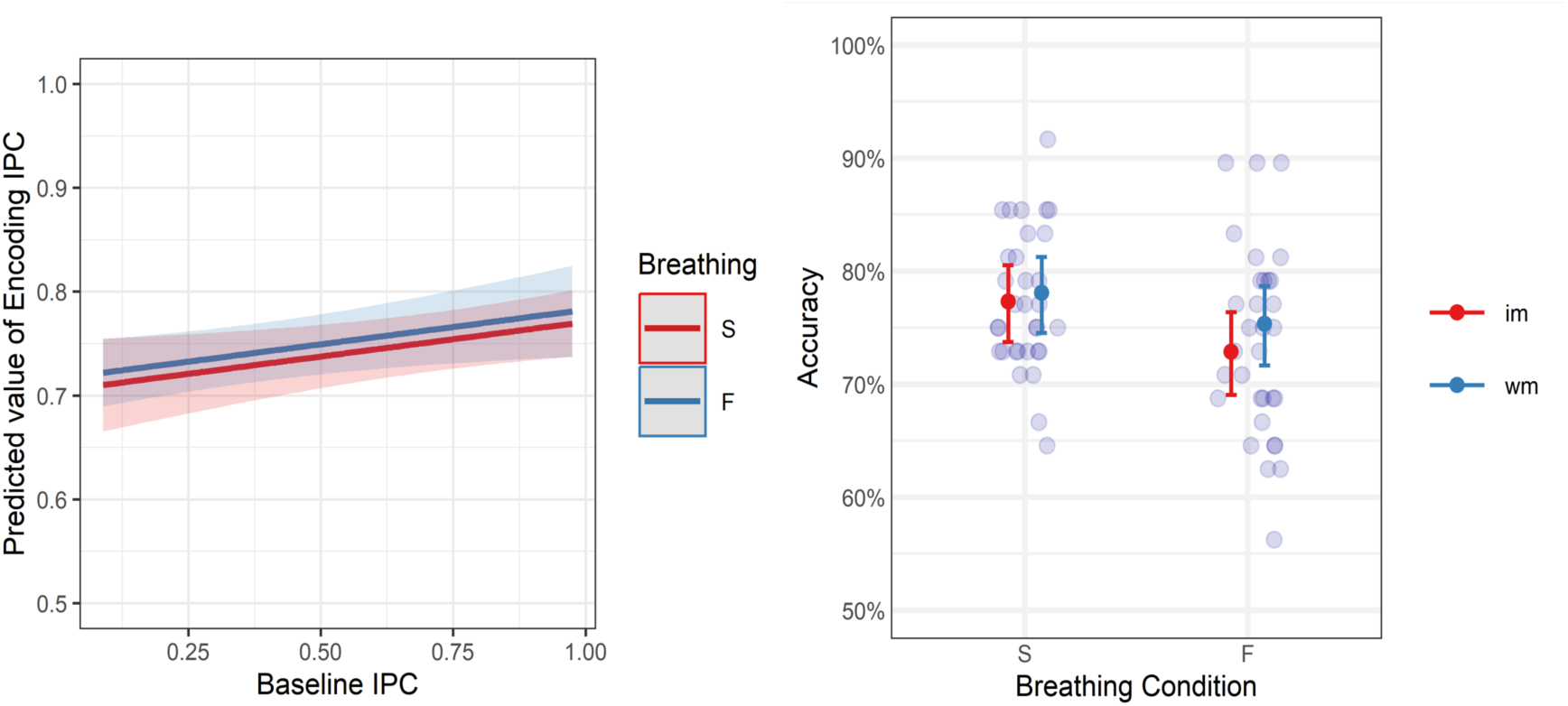
Persistence of enhanced coupling and behavioral effects. (A) Baseline cardiorespiratory coupling (measured during the final 60 seconds of paced breathing) predicted encoding-phase coupling during subsequent cognitive trials (b = 0.067, p = .020, d = 0.33), demonstrating that the coordinated autonomic state persisted even after breathing returned to natural rates. Points represent individual trials; lines show linear regression with 95% confidence intervals. Red = slow breathing; blue = fast breathing. (B) Slow breathing (6 bpm) improved behavioral accuracy compared to fast breathing (20 bpm) with equivalent effects across mental imagery (im) and working memory (wm) tasks. Error bars show mean ± SEM.

Consistent with coupling’s predictive relationship with accuracy, this experimental enhancement of coupling via slow breathing improved behavioral performance. Participants performed more accurately during slow breathing (M = 77.3%, SD = 6.1%) compared to fast breathing (M = 73.8%, SD = 8.3%), representing a 3.5 percentage point improvement (b = 0.238, SE = 0.118, z = 2.011, p = .044, OR = 1.27, R²m = 0.004, R²c = 0.026; Fig. 6B). This behavioral benefit was observed equivalently across both mental imagery and working memory tasks, supporting coupling as a domain-general mechanism.

Critically, respiratory frequency during task performance returned to natural rates in both conditions (slow breathing: M = 23.6 bpm, SD = 0.974; fast breathing: M = 23.4 bpm, SD = 1.060; t(30) = -0.456, p = .652), confirming no difference in breathing rate during cognitive trials despite the manipulation. This demonstrates that performance improvements resulted from the persisting coordinated autonomic state established during slow breathing, not from altered respiratory mechanics during task execution itself.

Importantly, respiratory frequency during task performance returned to natural rates in both conditions (slow breathing: M = 23.6 bpm, SD = 0.974; fast breathing: M = 23.4 bpm, SD = 1.060; t(30) = -0.456, p = .652), confirming no difference in breathing rate during cognitive trials despite the manipulation (slow pacing: 6 bpm; fast pacing: 20 bpm). This confirms that performance effects reflect carryover from the paced breathing manipulation rather than ongoing breathing rate differences during task execution.

Together, these findings establish the complete mechanistic pathway: slow breathing enhanced parasympathetic tone (RSA), which strengthened baseline cardiorespiratory coupling; this coordinated state persisted into cognitive trials even after breathing normalized, predicting encoding-phase coupling and improving behavioral accuracy.

## 4. Discussion

Cardiorespiratory coupling during the encoding phase predicted trial-level accuracy, whereas coupling during the maintenance phase showed no relationship with performance; this pattern was present for both the imagery and working memory tasks. This temporal dissociation indicates that synchronized cardiac and respiratory rhythms selectively predict performance when the brain actively constructs internal representations under elevated processing demands, not during passive retention of already-established representations. This temporal specificity constrains potential mechanisms underlying the coupling-cognition relationship.

The functional relationship between cardiorespiratory coupling and cognitive performance was supported through experimental manipulation of breathing rate. Slow breathing enhanced parasympathetic tone, strengthened baseline coupling, and improved behavioral accuracy, demonstrating that coupling strength can be manipulated to influence performance. Critically, while slow breathing enhanced both parasympathetic indices (RSA, RMSSD) and cardiorespiratory coupling, only RSA (which specifically indexes respiratory-linked heart rate variability) predicted baseline coupling strength, whereas RMSSD (a general measure of beat-to-beat variability) did not. This selective relationship reflects the physiological mechanism whereby vagal pathways modulate heart rate in synchrony with breathing, creating coordination between cardiac and respiratory oscillators in the brainstem (Schäfer et al., 1998); stronger parasympathetic tone during slow breathing amplifies this coordination.

Moreover, only encoding-phase coupling directly predicted trial-level accuracy; this relationship remained significant even when controlling for both RSA and RMSSD simultaneously, indicating that cardiorespiratory synchronization predicts performance independently of overall vagal tone. This dissociation extends the HRV-cognition literature, which has focused on parasympathetic activity as a unitary construct: parasympathetic enhancement facilitates coordination between cardiac and respiratory oscillators, but it is this synchronization—rather than vagal tone itself—that predicts cognitive performance. More generally, these findings suggest that cognitive performance may depend on the coordination of central and peripheral physiological systems, particularly during periods of elevated processing demand.

A dissociation emerged between objective performance and subjective awareness: coupling predicted trial-level accuracy but not vividness or confidence ratings. This indicates that cardiorespiratory coordination influences the actual precision and stability of internal representations without affecting metacognitive monitoring of representation quality. In other words, cardiorespiratory coordination may optimize physiological processes supporting representation construction without engaging higher-order metacognitive monitoring systems. This dissociation may have practical implications: individuals may not subjectively experience improvements from breathing interventions even when objective performance benefits occur.

Cardiorespiratory coupling predicted performance equivalently across mental imagery and working memory tasks despite their distinct encoding demands (verbal cue-based generation versus direct visual presentation). This generalization indicates coupling predicts the construction of internal representations regardless of input modality, suggesting a domain-general relationship rather than modality-specific optimization. The shared requirement across both tasks—transforming incoming information into stable internal representations maintained in working memory—appears to be the computational demand that benefits from cardiorespiratory coordination.

Respiratory frequency returned to natural rates (∼23 breaths/min) during cognitive tasks, yet coupling strength and performance benefits persisted. This demonstrates that slow breathing enhances cognition by establishing a coordinated autonomic state that outlasts the manipulation, rather than through direct respiratory mechanical effects during task execution. Baseline coupling predicted encoding-phase coupling even after breathing normalized. This persistence (∼12-minute task block) suggests breathing interventions could use brief preparatory protocols before cognitively demanding activities rather than requiring sustained altered breathing during performance. From a clinical perspective, this makes interventions more practical: patients could use brief preparatory breathing before exposure therapy or trauma narrative construction rather than maintaining altered breathing throughout treatment sessions. The duration and potential cumulative effects of repeated practice remain to be determined.

The selective benefit of cardiorespiratory coupling during encoding but not maintenance constrains potential mechanisms. Three non-exclusive pathways could account for this temporal specificity: 1) *Metabolic optimization:* Coordinated cardiac-respiratory rhythms may stabilize cerebral perfusion precisely when encoding imposes elevated oxygen demands, whereas maintenance of already-established representations requires minimal metabolic support (Willie et al., 2014). This account predicts coupling effects should be strongest during tasks with high metabolic requirements. 2) *Neural entrainment:* Synchronized peripheral rhythms could entrain cortical oscillations that facilitate binding distributed representations during encoding (Zelano et al., 2016), with less impact once representations stabilize. This predicts coupling should modulate neural oscillations specifically during encoding windows. 3) *Interoceptive signaling:* Stronger interoceptive coherence during high coupling states may enhance predictive processing when the brain must actively construct internal models. Arguably, all three accounts predict encoding-specific effects rather than domain-general performance enhancement, consistent with our findings. Distinguishing these mechanistic pathways definitively requires direct neural measurements—such as cerebral blood flow monitoring, EEG/MEG recordings during coupled breathing states, or fMRI examining network interactions during encoding—which remain for future investigation.

The 3.5% accuracy improvement from a single five-minute breathing session compares favorably to established cognitive interventions. HRV biofeedback training produces 3-8% improvements (Prinsloo et al., 2011; Lehrer & Gevirtz, 2014), while pharmacological cognitive enhancers yield 5-10% effects in healthy populations. Critically, our participants received no training or practice; sustained interventions report substantially larger benefits (Laborde et al., 2022). The effect generalized across both mental imagery and working memory tasks, suggesting broad applicability. For clinical populations with disrupted baseline coupling (anxiety disorders, PTSD, autonomic dysregulation), breathing-based interventions targeting cardiorespiratory coordination could potentially yield larger therapeutic benefits, though this requires empirical validation.

Our findings contribute to understanding mechanisms underlying breathing interventions’ therapeutic efficacy. Slow-paced breathing demonstrates effectiveness for anxiety and stress-related disorders (Laborde et al., 2022), with benefits typically attributed to enhanced parasympathetic tone indexed by heart rate variability. We confirm that slow breathing robustly increases parasympathetic activity (RSA and RMSSD), but additionally identify cardiorespiratory coordination as a parallel physiological marker predicting cognitive benefits. Clinically, this suggests interventions could monitor not only HRV but cardiorespiratory coupling strength to optimize outcomes. The specific temporal window when coupling predicts performance (during effortful encoding rather than passive processing) further suggests interventions could be strategically timed before cognitively demanding activities (e.g., exposure therapy, trauma narrative construction) to potentially maximize benefit, though controlled clinical trials are needed.

Several constraints on generalization should be acknowledged. First, we examined only visuospatial tasks; whether coupling predicts verbal memory, executive function, or semantic encoding remains untested. Second, slow breathing affects multiple co-varying systems (cerebral blood flow, baroreflex sensitivity, CO₂ levels), making it difficult to conclusively isolate coupling’s contribution from other autonomic changes. Third, while our correlational and experimental evidence converge, the trial-level relationship could reflect reverse causality if successful encoding inherently stabilizes autonomic rhythms; future studies using real-time coupling feedback or targeted disruption could address this. Fourth, we demonstrated state persistence across relatively short time-scales (the duration of the cognitive tasks); optimal intervention duration and whether effects accumulate with repeated practice require further investigation. Finally, and most importantly, without concurrent neural recordings we cannot determine the specific mechanisms by which coupling predicts encoding success. Our data establish *when* coupling matters but not definitively *how* it operates.

In conclusion, cardiorespiratory coupling predicts cognitive performance during encoding but not maintenance, establishing synchronized bodily rhythms as a physiological state marker for successful mental representation. Experimental enhancement through slow breathing improved accuracy, validating coupling as a functionally relevant and modifiable physiological parameter. These findings challenge accounts that treat neural computation in isolation from bodily physiology, revealing that optimal cognition is associated with coordinated peripheral signals. Clinically, cardiorespiratory coordination provides a specific, measurable target for optimizing breathing-based interventions, though the mechanisms linking coupling to cognitive benefits require direct investigation with neural imaging techniques.

## SUPPLEMENTARY MATERIALS

### Data Quality Control and Exclusions

Physiological data underwent quality control to identify excessive noise or technical failures. Exclusion criteria were: (1) noise exceeding 30% of the 5-minute paced breathing window, or (2) noise exceeding 20% during the 24-trial task window. Based on these criteria, data were excluded from: Participant 216 (WM task, fast-breathing condition), Participant 219 (WM task, both breathing conditions), and Participant 233 (all IM physiological measures due to data loss from computer update). In total, 386 trials (12.6% of 3,072 total trials) were excluded, leaving 2,686 trials for physiological analysis. For solely behavioral analysis, all of the 3072 trials were included.

### Model Diagnostics and Robustness Checks

Model assumptions were verified using simulation-based diagnostics (DHARMa package; Hartig, 2022) with 3,000 simulations for binomial models and Anderson-Darling tests for linear mixed models. Binomial models met all assumptions (see Supplementary Table S1). Most linear mixed models met normality assumptions without concerns (RSA: A = 0.573, p = .134; RMSSD: A = 0.409, p = .340; RMSSD-baseline IPC: p = .118). Three models showed significant Anderson-Darling tests, likely due to large sample sizes (>2,000 observations), but all exhibited acceptable skewness and kurtosis (|skewness| < 1, kurtosis < 5). Robustness analyses confirmed stability of all reported effects (see Supplementary Table S2).

**Supplementary Table S1.** : DHARMa Diagnostics for Binomial Models

DHARMA Diagnostics for Binomial Models
| Model | K-S<br>Test<br>D | K-S<br>p | Overdispersion<br>D | Overdispersion<br>p | Zero-<br>inflation<br>ratio | Zero-<br>inflation<br>p | Outlier<br>s p |
| --- | --- | --- | --- | --- | --- | --- | --- |
| Breathing effect on accuracy | 0.012 | .781 | 1.001 | .997 | 1.001 | 1.000 | .465 |
| IPC-accuracy, encoding phase | 0.014 | .626 | 0.014 | .626 | 1.00 | .930 | .267 |
| IPC-accuracy, maintenance phase | 0.014 | .658 | 1.004 | .917 | 1.004 | .933 | .267 |
| IPC×phases-accuracy | 0.006 | .977 | 1.003 | .941 | 1.003 | .947 | .805 |

**Supplementary Table S2.** : Robustness Checks for Linear Mixed Models with Significant Normality Tests

| Model | Original<br>estimate (b) | Bootstrap<br>95% CI | Robust<br>LMM b | Robust<br>LMM t |
| --- | --- | --- | --- | --- |
| Breathing effect on IPC encoding phase | 0.023 | [0.009, 0.040] | 0.023 | 3.06 |
| Baseline IPC predicting encoding IPC | 0.075 | [0.006, 0.122] | 0.075 | 2.73 |
| RSA predicting baseline IPC | 0.059 | [0.006, 0.122] | 0.059 | 4.774 |

*Note: Bootstrapped confidence intervals were consistent with original estimates and robust models yielded comparable parameter estimates and significance levels in all cases.*

